# Towards a mathematical understanding of colonization resistance in multispecies microbial communities

**DOI:** 10.1101/2021.01.17.426995

**Authors:** Erida Gjini, Sten Madec

**Affiliations:** Center for Computational and Stochastic Mathematics, Instituto Superior Tecnico, University of Lisbon, Lisbon, Portugal; Institut Denis Poisson, University of Tours, Tours, France

## Abstract

Microbial community composition and dynamics are key to health and disease. Explaining the forces generating and shaping diversity in the microbial consortia making up our body’s defenses is a major aim of current research in microbiology. For this, tractable models are needed, that bridge the gap between observations of patterns and underlying mechanisms. While most microbial dynamics models are based on the Lotka-Volterra framework, we still do not have an analytic quantity for colonization resistance, by which a microbial system’s fitness as a whole can be understood. Here, inspired by an epidemiological perspective, we propose a rather general modeling framework whereby colonization resistance can be clearly mathematically defined and studied. In our model, *N* similar species interact with each other through a co-colonization interaction network encompassing pairwise competition and cooperation, abstractly mirroring how organisms effectively modify their micro-scale environment in relation to others. This formulation relies on a generic notion of shared resources between members of a consortium, yielding explicit frequency-dependent dynamics among *N* species, in the form of a replicator equation, and offering a precise definition of colonization resistance. We demonstrate that colonization resistance arises and evolves naturally in a multispecies system as a collective quadratic term in a replicator equation, describing dynamic mean invasion fitness. Each pairwise invasion growth rate between two ecological partners, 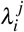, is derived explicitly from species asymmetries and mean traits. This makes the systemic colonization resistance 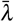 also an emergent function of global mean-field parameters and trait variation architecture, weighted by the evolving relative abundances among species. In particular, if the underlying invasion fitness matrix Λ displays *species-specific* ‘invasiveness’ or ‘invasibility’ structure, colonization resistance will be insensitive to mean micro-scale cooperation or competition. However, in general, colonization resistance depends on and may undergo critical transitions with changes in mean ‘environment’, e.g. cooperation and growth level in a community. We illustrate several key links between our proposed measure of colonization resistance and invader success, including sensitivity to timing, and to the intrinsic pairwise invasion architecture of the resident community. Our simulations reveal that *symmetric* and *invader-driven* mutual invasion among resident species tend to maximize systemic colonization resistance to outsiders, when compared to *resident-driven, anti-symmetric, almost anti-symmetric* and *random* Λ structures. We contend this modeling approach is a powerful new avenue to study, test and validate interaction networks and invasion topologies in diverse microbial consortia, and quantify analytically their role in colonization resistance, system function, and invasibility.

## Introduction

The human microbiota plays a crucial role in health and disease. Recent studies are increasingly uncovering one of its clearest contributions: protection against invading pathogens, known as colonization resistance (1; 2). This protection is important when enteric bacterial pathogens challenge the gastrointestinal tract, but applies also in the nasopharynx, skin, and other environments which are populated by rich consortia of commensal microbes. The composition of gut microbiota affects several collective metabolic functions and feedbacks with the host’s immune system, leading to or sometimes impairing immune homeostasis. Disruption of healthy microbiota, for example through antibiotics, or cytotoxic chemotherapy, can result in loss of colonization resistance and increased susceptibility to pathogens (3; 4), as well as long-term shifts in microbial ecosystem composition (5) and other diseases (6). Conversely, reconstitution of normal microbiota (e.g. through faecal transplants) has been demonstrated to restore host protection, and help cure patients from recurring pathogenic infections (7). Such dynamic processes of loss and gain of colonization resistance are particularly important in the intestinal tract, making stability, diversity and resilience of our gut microbial ecosystem an intense topic of investigation (2; 8; 9; 10).

Despite the progress in experimentally revealing the beneficial roles of the *>* 100 species (trillions of microbes) populating the human intestine (11), a theoretical understanding of how colonization resistance emerges as a collective trait from a community network, is maintained as a dynamic process throughout lifetime, and reacts to various perturbations, remains elusive. Mathematical approaches so far have been based primarily on Lotka-Volterra models (12; 13; 14) to explore ecological dynamics and stability of intestinal microbiota. However, while the generalized Lotka–Volterra equations typically capture qualitatively many empirical outcomes, they have a more limited ability to quantitatively recapitulate or parametrize observed dynamics (15; 16; 17). Furthermore, a mathematical definition and understanding of colonization resistance is still lacking. Explicit frameworks are needed for new perspectives on bottom-up community assembly and prediction of perturbation effects on the system, such as antibiotic administration, diet, immune suppression or host interaction with other drugs and vaccines. To address this knowledge gap, here, we synthesize a new conceptual and mathematical perspective to describe colonization resistance at the level of a single host, developing an analogy with an epidemiological multi-strain system (18).

Our model is derived from a time-scale separation for coexistence dynamics between similar strains, which yields explicit *N*-dimensional replicator dynamics for their relative abundances, while conserving global carrying capacity (18; 19; 20). Despite being the central equation of evolutionary game theory (21), this equation is surprisingly little used in microbiology, in favour of the predominant use of generalized Lotka-Volterra (gLV) frameworks for total abundances (12; 14; 22). Although there are proven mathematical links between the two (23), these formulations are not exactly equivalent. In our setup, the frequencies of multiple species are central, as on one hand, only relative abundances between (pseudo-)species of interest are most often empirically accessible via metagenomic sequencing data, and on the other hand, in our model these can be fully analytically predicted via replicator dynamics in terms of the magnitudes of pairwise invasion fitnesses in the system 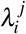. These fitness coefficients reflect initial demographic performance in dyadic *i* → *j* invasion scenarios between two species, starting from low frequency of *i* in an equilibrium set by *j*, and besides being potentially easier to measure, constitute a fundamental concept in the theory of adaptive dynamics (24), and in our case are entirely trait-mediated in a bottom-up fashion. Our approach allows thus to define a clear measure of colonization resistance, related to these replicator dynamics and 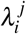, to explore how this measure affects outsider invasion performance across ecological networks, and over time within the same system, and to propose a new basis for fresh in-depth study of community assembly or response to perturbations.

Application of this model to in-host microbiota dynamics requires re-interpretation of original variables and parameters, and a conceptual shift of scales. In the following, we present the key ingredients of this modeling approach and develop the analogy for translation of the model to the within-host level. Finally, we illustrate several ways in which this model can be used to provide new insights and integrate several findings on microbial colonization resistance. Simulations are conducted in Matlab R2017b (25) and Python 3.10 (26).

## Results

### A host is a system of ‘free’ and ‘occupied’ micro-niches that can be colonized and co-colonized

We define the host as a well-mixed system, and the dimensionality of the system as the number of similar microbial types (entities) interacting in colonization of such a system, hereafter denoted as ‘species’. In the re-framing following (18), we model the within-host environment as a number of potential micro-’niches’ depicting generic growth units (resources) to be utilized, and we track the proportion of such niches that remain free or susceptible (*S*), those that get singly-colonized by either species (*I*_*i*_), and those that get co-colonized, either twice by the same species (*I*_*ii*_) or by two different species (*I*_*ij*_) (see Methods, and Figure 1A). By taking suitably small such growth units within-host, we can truncate the multiplicity of ‘infection’ (MOI) to at most 2 species per unit. It is assumed that such growth units (similar to hosts in the epidemiological setting) homogeneously mix in the system, and when they are free, they are equally accessible to all species, whereas when they are already colonized, they become more or less available to co-colonization/co-utilization by others, depending on the identities of the interacting partners.

**Figure 1.**
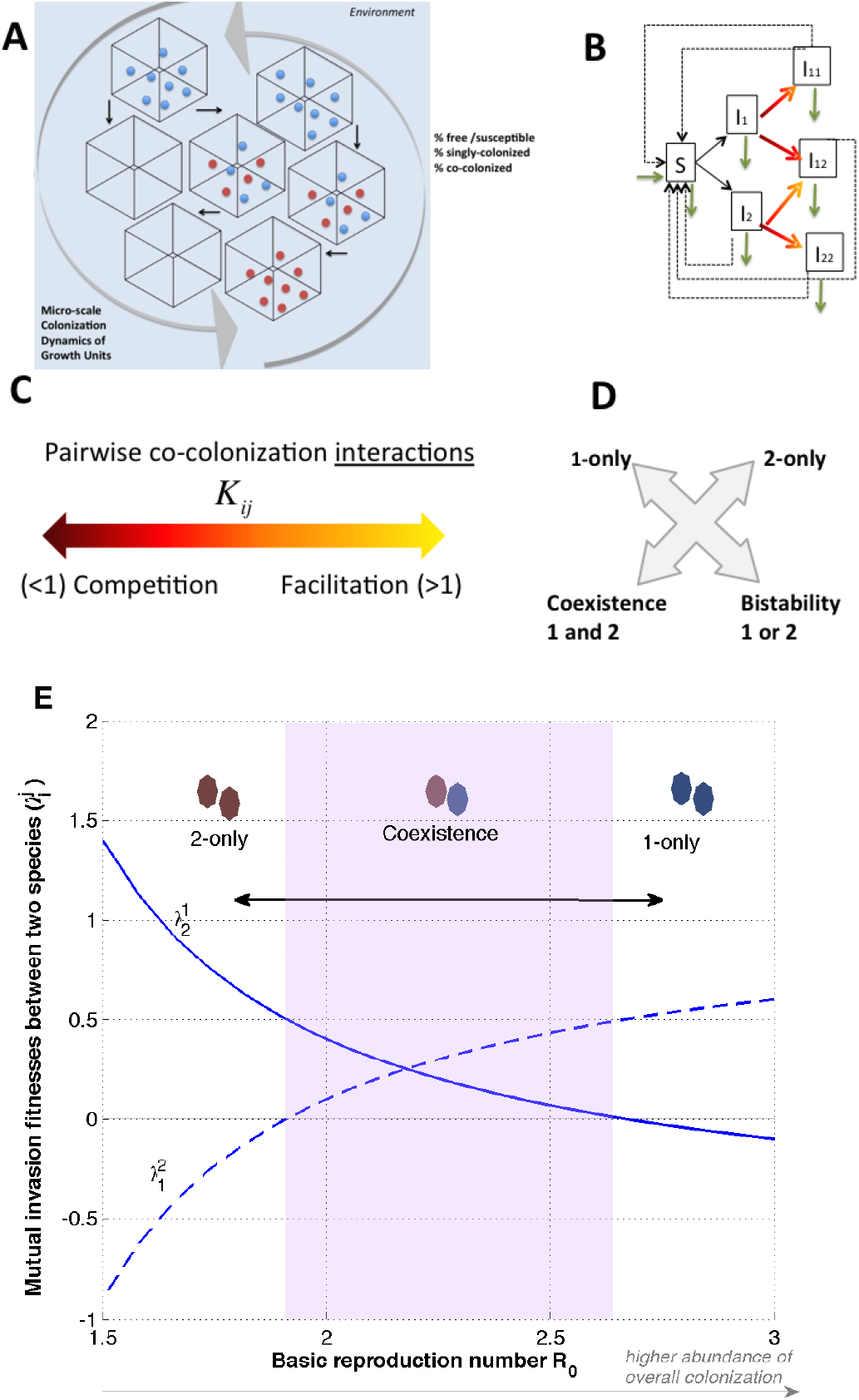
Analogy between epidemiological and within-host multi-species dynamics (here illustrated for *N* = 2). The model is based on SIS dynamics with co-infection (18). **A**. Conceptualization of the within-host environment, where micro-niches can be free (uncolonized *S*), singly- (*I*_*i*_) or doubly-colonized (*I*_*ii*_,*I*_*i j*_, *I*_*j j*_). Full equations are given and analyzed in (18). **B**. Model structure for niche state transitions, including colonization (black arrows), co-colonization (color-shaded arrows) and clearance (dashed arrows). Natural birth/death is denoted by green arrows. **C**. Singly-colonized patches may have a reduced or increased susceptibility to becoming co-colonized, noted by coefficients *K*_*ij*_ (*> / <* 1) between any two species. **D**. For *N* = 2, there are four outcomes (Table 1), depending on species interactions: exclusion of either, coexistence, or bistability. **E**. Context-dependence of mutual invasion fitnesses 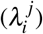 generates different ecology between 2 species (*K*_*ij*_ fixed). Starting from coexistence, lowering overall colonization (*R*_0_ ↓), e.g. via antibiotics, or increasing it drastically, may lead to competitive exclusion favouring an opposite species in each direction.

Within each single growth unit, microbial propagules can grow, then upon release, can be transmitted to other units. Microbial propagation dynamics, thus, under homogeneous mixing of propagules in the in-host milieu, can be described, to a first-order approximation through simple mass-action kinetics (Figure 1B). The growth of each microbial species depends on their transmission rate *β* and frequency of niches ‘emitting’ propagules of that species. The transmission parameter, *β*, in this analogy, can be taken as a net parameter, encapsulating the chain of events including local growth happening in the micro-niche of ‘origin’, followed by migration, and diffusion of infectious propagules in the system, multiplied by the effective probability of settling in other ‘destination’ niches. In its basic version, the model assumes *β* is equal for all *i*, yielding reproductive fairness in the system (but see (20) for a formal proof and generalization to a higher number of traits varying among relatively similar species).

Interactions among species happen upon co-colonization, where the colonizer (*i*) and co-colonizer (*j*) microbes may compete or facilitate each other, each ordered pair with its own coefficient *K*_*ij*_ (Fig. 1C). Even without specifying the explicit ecological mechanisms, which may be synergistic or antagonistic, and include cross-feeding, antagonistic warfare via bacteriocins, local nutrient alteration or depletion, acidification of the micro-environment, substrate availability, and others (27; 28; 29; 30; 31; 32), this implies that what counts at an abstract level is that the micro-scale presence of one species alters the value of that same resource-growth unit complex for individuals of that same/other species, making it better or worse. Such altered ‘susceptibilities’ to co-colonization *K*_*ij*_ (*>* 1 : facilitation, *<* 1 : competition), for *N* = 2, form a 2-by-2 matrix (Figure 1C), which determines one of four possible outcomes between 2 species (33): exclusion of 1, exclusion of 2, coexistence or bistability (Fig. 1D). However, the cocolonization interaction matrix in general is *N*×*N*, when considering *N* possible ecological partners in the community. Singly- and co-colonized units are assumed to produce an equal number of propagules, and mixed co-colonization units *I*_*ij*_ produce and transmit *i*/ *j* propagules with unbiased probability 1*/*2.

Free units of resource are replenished at rate *r* (in-flow), assumed equal to the natural decay rate (out-flow), keeping thus constant the carrying capacity of the system. Within each available micro-niche microbes can grow for a mean duration of 1*/γ* time units, hence yielding a turnover rate of *m* = *r* + *γ*. The basic reproduction number *R*_0_, denotes how many new colonizations, each colonized micro-niche will produce over its lifetime as occupied, if occurring in a totally colonization-free environment, and is given by *β/m*, similar to epidemic models: if *R*_0_ *>* 1 there is a globally stable endemic state with 1− 1*/R*_0_ colonization prevalence. If *R*_0_ *<* 1, the system is colonization-free. With this established analogy, the *N*-species model (18) can be applied to microbiota dynamics within host, as a first-order approximation, in much the same spirit as a Lotka-Volterra dynamic model, but here with the unit of resource, whether *free* or *shared*, made explicit. Because our model does not have a spatial extent, like most gLV models applied in microbial ecology (22), it can be considered as similar to the single-chamber, continuous flow chemostat, which is often an *in-vitro* model system used to study multi-species asssemblages and dynamics (34; 35; 36; 37; 38), albeit possibly with more complex substrate interaction and turnover dynamics, or multiple spatial compartments, compared to the one assumed here. For a more explicit derivation of our analogy via first-principles and a consumer-resource description, please refer also to Supplementary Table S1.

### Explicit in-host frequency dynamics of *N* species under homogeneous mixing

Considering phenotypic similarity in micro-scale interaction trait space between species (*K*_*i j*_ = *k* + *εα*_*i j*_, with *ε* small), we have shown for *N* = 2 and for general *N* ≥2 in (18), that there are two timescales in such a system: a *fast* one where the species obey neutral dynamics, during which global colonization and co-colonization variables stabilize, and a *slow* timescale, where non-neutral dynamics happen, driven explicitly by species variation in the co-colonization interaction matrix *K* (or more traits as shown in (20; 39)). Applying the similarity assumption in *N* dimensions (18) (where *k* can reflect the mean of *K*_*ij*_, and *ε* their standard deviation), corresponds to a kind of Taylor approximation for the dynamics, which gets decomposed into a sum of different order terms. Thus, we have mapped the *N*-species dynamics explicitly to a special replicator equation (21) for the change in species frequencies over the long timescale *τ* = *εt*:

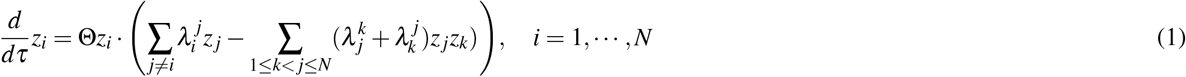

with Σ_*i*_*z*_*i*_ = 1 and 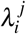 denoting mutual invasion fitnesses between species (Table 1) and Θ the speed of selection. The mutual invasion fitness, 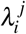, is a key quantity at the heart of adaptive dynamics (24), and depicts the initial demographic performance, namely the invasion growth rate of a species *i* in an equilibrium set by a single resident *j*. It is typically derived from the underlying ecology of the model. In our model, with variation only in co-colonization coefficients, this pairwise invasion fitness is given by:

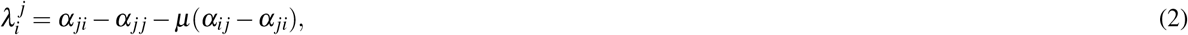

and is a direct function of the phenotypic variation in co-colonization interactions (the *α*_*i j*_’s), and of the ratio of single to co-colonization in the system, *µ*. Notice that for any entity *i* in the system, the rate of growth of its frequency (*z*_*i*_) relative to other entities, is given by 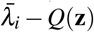, namely by how it performs in pairwise invasion ‘games’ with any other member of the system (with 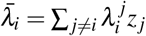, summed over all *j*), and also by how the entire system as a whole (*Q*) acts to promote or reduce net individual growth. The dynamic quadratic term *Q* in this replicator equation equals precisely mean invasion fitness, hence colonization resistance in the system:

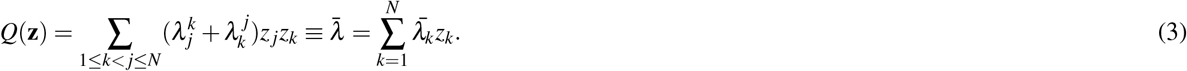

**Table 1.** Key features of the co-colonization model for similar species (18)

| The constituent variables and parameters are the following: |  |
| --- | --- |
| Micro-scale co-colonization interactions ($N \times N$) | $K_{ij} = k + \epsilon \alpha_{ij}$ (similarity assumption: $0 \leq \epsilon \ll 1$) |
| Normalized interaction strengths | $\alpha_{ij} = \frac{K_{ij} - k}{\epsilon}$; (e.g. $k$ = mean of $K_{ij}$) |
| Basic reproduction number | $R_0$ |
| Equilibrium colonization prevalence ($I^* + D^*$) | $1 - S^* = 1 - 1/R_0$ |
| Ratio of single to co-colonization ($I^*/D^*$) | $\mu = [(R_0 - 1)k]^{-1}$ |
| Speed of multispecies dynamics | $\Theta = \beta \left(1 - \frac{1}{R_0}\right) \left(\frac{\mu}{2(\mu+1)^2 - \mu}\right)$ |
| Timescale of selection (slow timescale) | $\tau = \epsilon t$ |
| Frequency of species $i$ | $z_i(\tau)$, with $\sum z_i = 1, i = 1, \dots, N$ |
| Single and co-colonization prevalence $i/(i, j)$ | $I_i = I^* z_i; I_{ij} = D^* z_i z_j$ |
| Initial species composition and rel.abundances | $\mathbf{z}(0)$ |
| Trait-based mutual invasion fitness, in this model*: |  |
| Invasion growth rate of $i$ in a $j$-only equilibrium | $\lambda_i^j = \alpha_{ji} - \alpha_{jj} - \mu(\alpha_{ij} - \alpha_{ji})$ |
| Frequency dynamics for $N = 2$ are given by: | |
| $\frac{d}{d\tau} z_2 = \Theta z_2 \cdot \left[ \lambda_2^1 (1 - z_2) - (\lambda_2^1 + \lambda_1^2) z_2 (1 - z_2) \right], z_1 = 1 - z_2$ | |
| $(N = 2)$ 4 mutual invasion structures | 4 outcomes: types of edges ($N$ species network) |
| $(\lambda_1^2, \lambda_2^1) = (+, -)$ $1 \leftarrow 2$ | 1 wins in competitive exclusion: $z_1^* = 1, z_2^* = 0$ |
| $(\lambda_1^2, \lambda_2^1) = (-, +)$ $1 \rightarrow 2$ | 2 wins in competitive exclusion: $z_1^* = 0, z_2^* = 1$ |
| $(\lambda_1^2, \lambda_2^1) = (+, +)$ $1 \text{ --- } 2$ | 1 and 2 coexist: $z_1^* = \lambda_1^2 / (\lambda_2^1 + \lambda_1^2), z_2^* = 1 - z_1^*$ |
| $(\lambda_1^2, \lambda_2^1) = (-, -)$ $1 \text{ --- } 2$ | Bistability: $z_1^* = 1$, or $z_2^* = 1$ depending on $z_1(0)$ |
| Environmental scenarios | Effect on parameters ($\dots \rightarrow \Lambda, \Theta$) |
| Factors promoting colonization for all species | $R_0 \uparrow$ |
| Increased microbial antagonism (or facilitation) | $k \downarrow$ (or $k \uparrow$) |
| Different species number/composition | Different $N/K_{ij}$ entries |
| Metabolic cross-feeding/bacteriocin effects | Possibly different $K_{ij}$, different $R_0$ |
| Broad spectrum antibiotic treatment $\uparrow (\downarrow)$ | $R_0 \downarrow (R_0 \uparrow)$ |
| Different nutrient substrate/growth efficiency | Different $\beta$ |
| The system with $N$ interacting species evolves according to replicator dynamics: | |
| $\frac{d}{d\tau} z_i = \Theta z_i \cdot \left( \sum_{j \neq i} \lambda_i^j z_j - \sum_{1 \leq k \neq j \leq N} \lambda_j^k z_j z_k \right), \quad i = 1, \dots, N$ | |
| Colonization resistance as mean invasion fitness of the system |  |
| $Q(\mathbf{z}) = \sum \sum_{1 \leq k \neq j \leq N} \lambda_j^k z_j z_k = \sum_j \bar{\lambda}_j = \bar{\lambda}$ | |
| $Q$ captures the tension between individual and group success in the system, as species dynamically shape their common environment. | |
| If $Q > 0$, the system as a whole acts negatively on each species, making $Q$ a cost to any current member of the consortium, but by consequence, affecting negatively also invaders, thus a net benefit for the external stability of the system.<br>If $Q < 0$, the system as a whole acts positively on all species, increasing their growth rate, but also facilitating invaders.<br>$Q$ can be seen as a function of time ($\tau$), of species composition and relative abundances ($\mathbf{z}$), and of pairwise invasion coefficients ($\Lambda$) |  |

\* But see (20; 39) for a more general derivation in the case of 5-trait dimensions where similar species can vary.

It sums the global effect of the system on each member, and reflects mathematically whether the multi-species consortium as a whole adds (*Q <* 0) or detracts from the growth of each individual member (*Q >* 0) (18).

More generally, in a scenario when more traits are allowed to vary between species, the trait-based pairwise invasion fitness expression is more complex, but the same replicator dynamics can be derived (20). Whatever their exact expression, together the 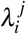 between all pairs, drive collective dynamics in the system. It is clear from equation 1, that to survive in the system, an individual/species must always ‘play games’ with co-occurring partners, and the payoff depends not only on an individual’s own (vector) strategy 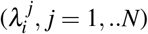, but also on strategies and abundances of its opponents. Because *Q* involves a product of frequencies, it is non-zero, only when there are more than 1 species coexisting, and when 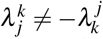 for all pairs (*j, k*). In the simplest system with only two species, *N* = 2 (Table 1), it is easy to see that at the coexistence steady state, *Q*(*z*_1_, *z*_2_) is positive and maximized for *z*_1_ = *z*_2_ = 1*/*2. Thus, a 2-species system, on its own, independently of how a third invader interacts with each member separately, is mostly protected against such invader, when its two constituent members are equiabundant (see Fig. 2).

**Figure 2.**
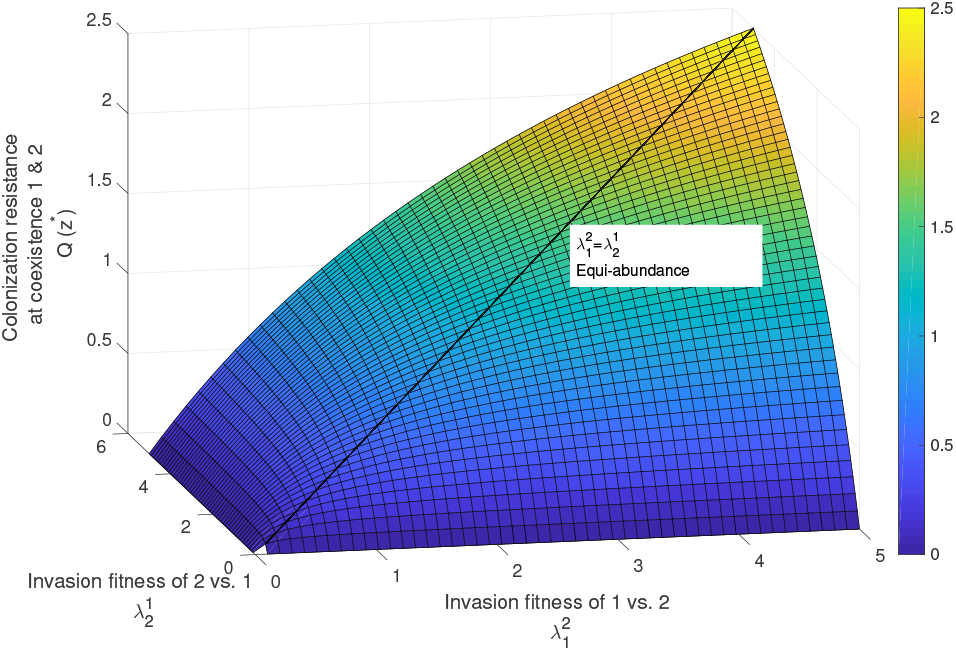
Colonization resistance *Q* in the *N* = 2 system with 2 species coexisting. Here we visualize a 3-dimensional plot of *Q*(*x, y*) = *xy/*(*x* + *y*) as a function of mutual invasion fitnesses 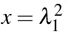 and 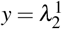 when the system is at the coexistence equilibrium with species frequencies 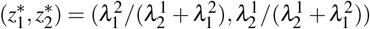. The maximum of *Q* is reached at 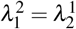, i.e. for equiabundance of two resident species, and it increases with the absolute magnitude of the *λ* ′s.

### Colonization resistance: system vs. outsider species

Depending on how many species make up the resident species pool, *N*, and how they interact, one models a different system size and a different network of mutual invasion fitnesses (directed edges with precise magnitudes). This will generate system-specific frequency dynamics eventually leading up to a pruned community with *n*≤ *N* species, and simultaneously the evolution of system resistance to invasion by outsiders (Table 1). The micro-scale effects between species upon co-colonization are captured by the *K*_*ij*_, denoting a fixed microscopic pairwise interaction trait, which, if above 1, denotes facilitation between *i* and *j*, and if below 1, denotes competition. The mutual invasion fitnesses between species, on the other hand, 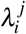 (positive or negative), emerge as higher-level and context-dependent traits (40), in this case determined by the global basic reproduction number *R*_0_ and mean interaction coefficient *k*, (but more generally may depend on additional mean-field parameters as shown by (20)). What is key in 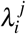 is that the invasion performance of *i* into a *j* − *only* equilibrium, depends on how much the resident entity *j* relatively favours the invader *i* over itself (trait difference *α*_*ji*_ − *α*_*jj*_ in 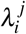) and also on the relative benefit of *i* from transitions to mixed co-colonization with *j* (−*µ*(*α*_*i j*_ − *α*_*ji*_) term). This second term, depends on relative availability of singly-colonized micro-niches to co-colonize in the system, hence on the ratio *µ* = *I/D* = 1*/*((*R*_0_ − 1)*k*). Thus there are two effects in 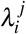:*i*) comparative advantage from non-self to self interaction of the resident, and this is not dependent on co-colonization prevalence in the system; and ii) relative benefit from asymmetric investment in mixed co-colonization, and this decreases with prevalence of co-colonization in the system. Notice that whether the species are competing or cooperating in their use of resources (*k* small tending to zero, or *k* large tending to a number much higher than 1) has effect only on the second term of 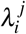 via the ratio *µ*, if *α*_*ji*_ ≠ *α*_*i j*_, hence only for non-symmetric co-colonization interactions *K*_*ij*_.

It is the 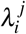 that ultimately govern relative species abundances, co-occurrence, diversity, stability, and system properties (see Supplementary Movies S1-S2). Very crucially, pairwise *i* − *j* coexistence for 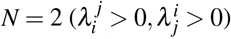 does not imply *i* − *j* coexistence in a community with more species, because in the network with more species, how these two members interact individually with third-parties, alters their net growth rate and ultimately their fate in the system.

### Special types of mutual invasion networks between N species

In our previous work (18), we have sketched several canonical mutual invasion structures for the matrix of 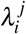 in a system with *N* members, i.e. invader-driven (equal columns); resident-driven (equal rows); symmetric (Λ = Λ^*T*^); anti-symmetric (Λ = −Λ^*T*^); and random, and outlined some corresponding feasible structures of the co-colonization interaction matrix with *K*_*i j*_. For example, symmetric *K*, such that *K*_*i j*_ = *K*_*ji*_ = *K*_*ii*_ + *K*_*jj*_ − *k* leads to invader-driven Λ; whereas symmetric *K*, such that *K*_*i j*_ = *k* for *i* ≠ *j*, with the diagonal entries free to vary, leads to resident-driven Λ. (see Table 2 in (18) for more details). However, there are in principle infinitely many possible microscopic *K*_*ij*_ leading to the same macroscopic 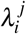 architecture. Here, we revisit these canonical structures in the context of their implications for colonization resistance of a completely closed microbial ecosystem. Note that in some cases, the net 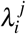 may be easier to access and measure empirically than the underlying micro-scale co-colonization interactions *K*_*ij*_. In even more complex scenarios, when more than one fundamental traits or biological processes vary between species (20), the problem of measuring several trait dimensions to obtain 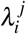 bottom-up, gets substantially reduced if one focuses just on quantifying the net pairwise invasion fitnesses in a top-down and appropriately coarse-grained manner.

For symmetric mutual invasion 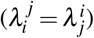, *Q* always increases (*dQ/dt >* 0), and this case corresponds to Fisher’s fundamental theorem (41). In this particular case, cycles among species are impossible, and their dynamics converge typically to stable coexistence fixed points, and competitive exclusion is unlikely. The implication of this monotonic scenario is that under perfectly symmetric invasion growth rates, when we observe the same system at different levels of colonization resistance, we can assign a temporal order to different observations, distinguishing which community resilience came first in time.

In the opposite extreme, for anti-symmetric mutual invasion 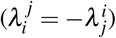, the system colonization resistance is exactly zero over all time, *Q* = 0, corresponding to zero-sum games in evolutionary game-theoretic terms (42). Dynamics in that case tend to a (structurally unstable) family of cycles around a center, like in Lotka-Volterra prey-predator models (43), where an odd number of species coexist. More special cases, such as the species-centric invader-driven invasion and resident-driven invasion, and explicit *K*− Λ matrix (trait - invasion) links are analyzed in (18).

Investigating community dynamics under these pairwise invasion architectures, we find that keeping all else equal, and starting from random initial frequencies, *Q* increases fastest over time in a system regulated by *invader-driven* mutual invasion (Fig. 3), i.e. where species are ‘defined’ by their active invasiveness trait. This case typically leads to coexistence among several species and rarely to exclusion. In the resident-driven mutual invasion case, where species are defined by how they allow invasion by others, hence by their invasibility, dynamics tend most likely to competitive exclusion, and hence toward *Q* = 0, but following an initial selection period during which systemic *Q* is negative.

**Figure 3.**
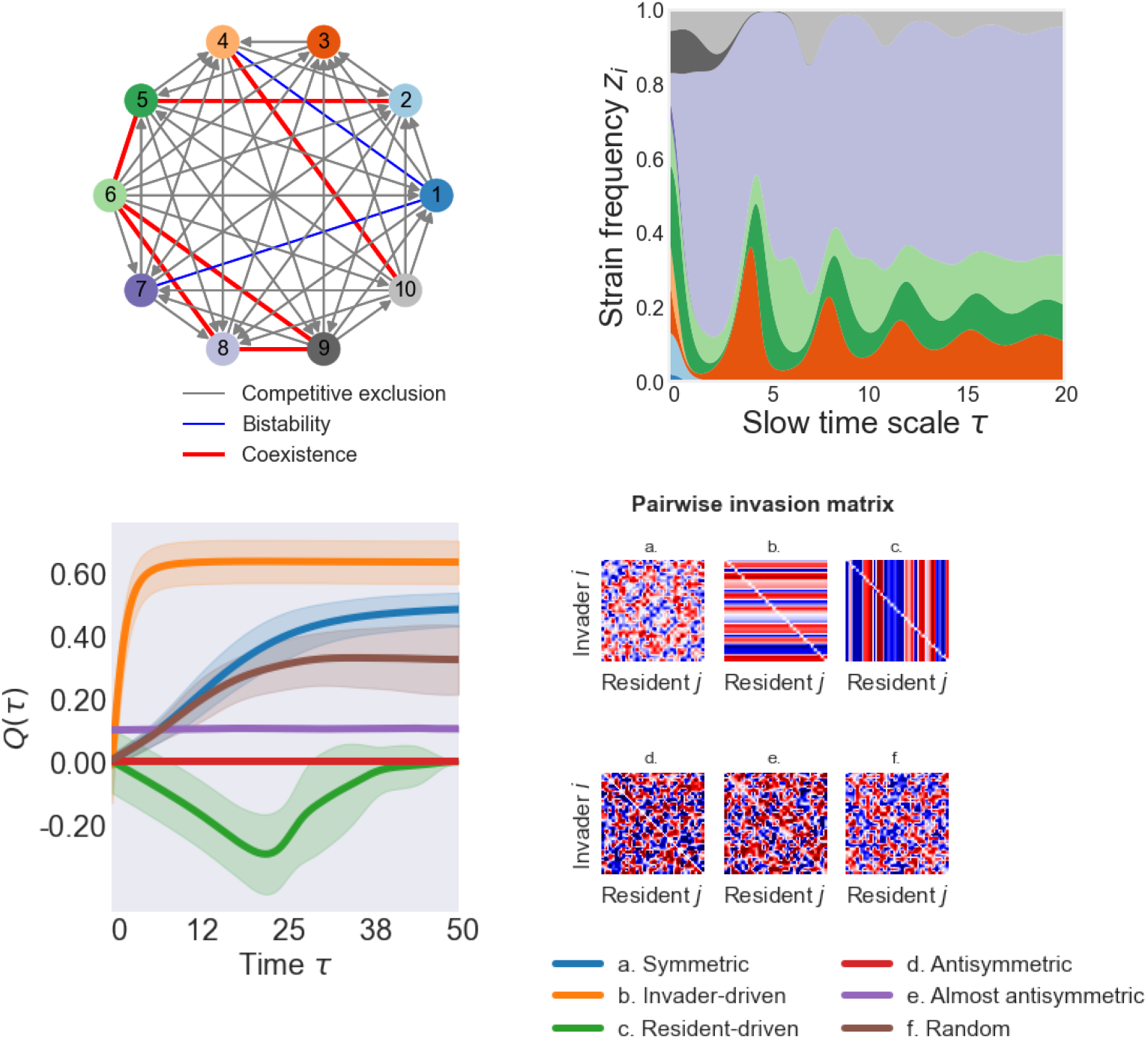
Mutual invasion structure of the microbial ecosystem, species dynamics and system mean invasion fitness. **Top panel:** Illustration of a mutual invasion network between 10 species, whose structure and pairwise invasion fitness values lead to multispecies dynamics on the right (Eq.1), with ultimately 5 species coexisting. **Bottom panel:** Colonization resistance for special invasion structures. On the left, we plot *Q* dynamics (mean: lines, ± sd.: shading) over 30 stochastic realizations for 6 cases of Λ between *N* = 30 species (shown on the right). For each case, random uniform 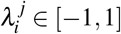, from the same distribution and range, are drawn, and fed each into Eqs.1 (assuming Θ = 1), which are then solved with random initial relative abundances *z*_*i*_(0), as in (18). *<u>Symmetric</u>* Λ yields dynamics described by Fisher’s fundamental theorem, where mean invasion fitness always increases and stable coexistence is likely. *<u>Invader-driven</u>* Λ implies large coexistence potential among multiple species. *<u>Resident-driven</u>* Λ implies large potential for competitive exclusion, and favours a single species becoming the only persistent one of the community. *<u>Antisymmetric</u>* Λ makes *Q* ≡ 0, corresponding to zero-sum games among species and complex structurally unstable coexistence dynamics. *<u>Almost-antisymmetric</u>* Λ typically produces complex but more regular dynamics, for example limit cycles, leading to periodic *Q. <u>Random</u>* mutual invasion allows rich multi-species dynamics to unfold. Cases **a-c** can be obtained for 3 particular instances of symmetric co-colonization interactions matrix *K* (18).

**Figure 4.**
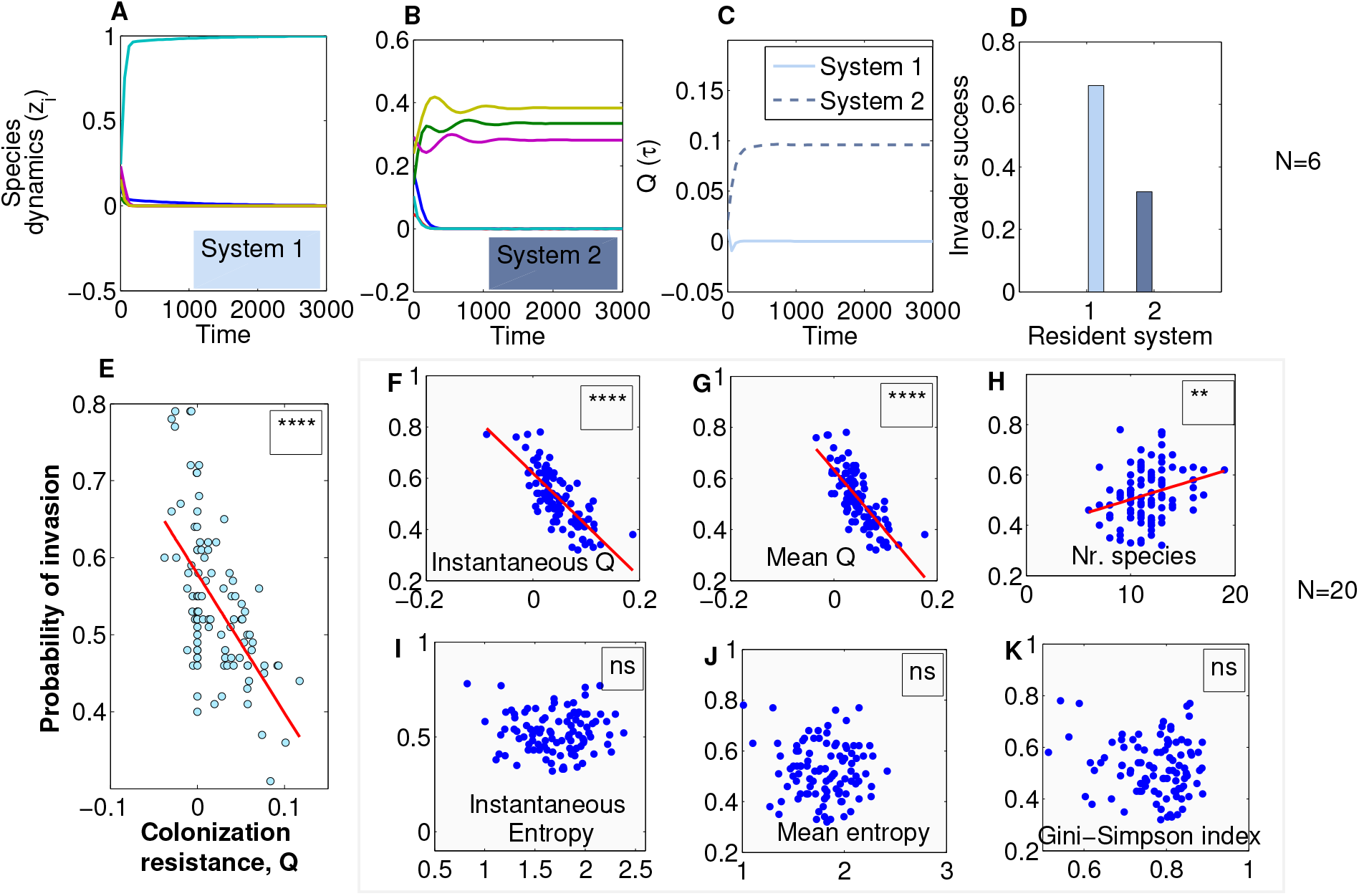
Colonization resistance, as defined by our model, and outsider invasion. In **A-D**, we fixed the multispecies *K*_*ij*_ interactions in susceptibilities to co-colonization in 2 systems with *N* = 6, *β* = 1, *k* = 0.3, *R*_0_ = 2. We simulated 100 realizations of invasion by an outsider species, sampling its interaction traits from the same distribution as the resident 𝒩(*k, ε*^2^), and introducing it at the mid-point of the interval. We counted invasion successful if the invader grew from its low initial frequency (here 10^−4^). System 2 (higher *Q*) was superior in preventing invasion to system 1. In **E**, we repeated the procedure for 100 randomly generated systems (blue dots) and examined system *Q* at invasion and invader success, finding a significant negative relationship between colonization resistance and invasibility (*p <* 10^−5^). The final outcome of invasion whether the invader ultimately coexists, or how original species are affected was not studied. In **F-K**, we examined invasion success for *N* = 20. Simulating 100 random systems as in **E**, we regressed outsider invasion probability on different system characteristics: *Q* (*p <* 10^−5^) in **(F-G)** and the number of species in the system (*p <* 0.01) in **(H)** were significant, while diversity indices **(I-K)** were not.

There is undoubtedly a vast complexity of possible co-colonization interaction networks and emergent invasion architectures between *N* species which yield coexistence scenarios of more complex nature than fixed points, including multistable attractors, limit cycles and chaos (19). Yet, in all these cases, the net colonization resistance that arises and changes as an adaptive function of an evolving ecosystem can be mapped directly to the relative frequencies of constituent species.

Thus, depending on how frequency dynamics unfold, subject to intrinsic or extrinsic drivers, *Q* dynamics can be higher or lower, monotonic or oscillatory, and feed back on the system, in a fully analytically-explicit manner (Table 1).

Remark that even when co-colonization interactions are symmetric in the system, i.e. the species pairs favour or inhibit each other in the same way for sharing a common resource unit, independent of the order of landing on that resource (*K*_*i j*_ = *K*_*ji*_), the effective mutual invasion fitness coefficients (the 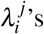) that emerge can be asymmetric (18) depending on further constraints upon that symmetry, leading ultimately to very distinct hierarchical dynamics (e.g. species-specific invader-driven Λ, or resident-driven fitnesses), with rather opposing effects on *Q*.

### Antibiotics can change colonization resistance and effect depends on species interactions

It is precisely the analytical tractability of this model (18; 19) that makes it easier to study global perturbation effects on the system, and to interpolate across conditions (Table 1). Perturbations affecting global context, such as *R*_0_ and *k*, directly change the mutual invasion fitnesses 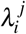, and thus species dynamics. An example are broad-spectrum antibiotics, leading to alteration of *R*_0_ in the system, likely reducing the colonization of all species, and shifting their internal competitive balance.

Even when acting symmetrically on system members, more antibiotics in our model can drive the system from coexistence to competitive exclusion (see the case of *N* = 2 in Fig. 1E.), hence toward *Q* = 0, promoting outsider invasion. Through our explicit 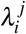 (18), one can clearly see that this *R*_0_ impact drastically varies, depending on the interaction network between species, for example it is stronger when *k* is lower, i.e. when competition is higher in the system (see Supplementary Movies S3-S4). In particular, even when starting at same baseline invasion fitnesses between two species (i.e. also the same coexistence scenario in terms of relative abundances), the stronger the mean competition in micro-scale co-colonization (the lower *k*), the more drastic will be the selection in the system induced by the antibiotic. Such explicit individualization of perturbation effects, based on underlying interactions, may be easily related to data and to context-dependence of outcomes in microbial consortia.

In more diverse communities (more species *N >* 2, and/or more traits varying), effects of context-altering perturbations are undoubtedly more complex, as we have shown in some cases (19; 20; 39). Among *N* species varying randomly only in co-colonization interaction phenotypes (susceptibilities *K*_*ij*_), a key modulator of coexistence regimes is the ratio of single-to co-colonization in the system, *µ* (19), decreasing with *R*_0_ and *k*. We have shown that in this model, *µ* = 1*/*((*R*_0_−1)*k*) acts as an axis of tunable contextual feedback on the multispecies dynamics: for standard normally-distributed rescaled interaction strengths (*α*_*i j*_), when *µ*→ ∞, and all else is kept fixed, the mutual invasibility network tends to contain mostly competitive exclusion edges, which makes more species coexist, but in unstable and oscillatory fashion, contrasting the regime of *µ*→ 0 displaying mostly stable steady states (19). Thus, decreasing *R*_0_ in such a scenario, leading possibly to a fluctuating *Q* via lower colonization prevalence and higher *µ*, can make a given system more vulnerable to opportunistic invasion, particularly when *Q* hits low or negative values, even if transiently. This seems to mimick what has been observed empirically, for example in individuals who following antibiotic therapy experienced domination of their gut microbiota by pathogenic *E. faecalis* or *C*.*difficile* bacteria, and succumbed to infection (3; 44). Notice that in our model, a similar effect could be obtained by decreasing *k*, hence decreasing cooperation, in the multispecies system: the ratio of single-to-cocolonization *µ* would increase, thereby promoting lower and oscillatory colonization resistance via fluctuating multispecies dynamics (19).

### Invasion by outsiders depends on colonization resistance

Mathematically, it is straightforward to see that the initial growth rate of any invader in a system at state **z**(*τ*) is given by:

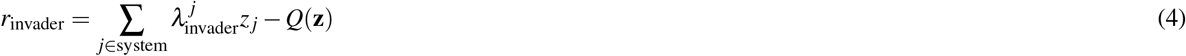

The first summation term in *r*_invader_ depends on invader traits, 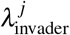, namely how this species invades any existing species currently in the system. The second term instead, −*Q*, is independent of the invader, and describes how the system colonization resistance by itself, detracts from invader’s initial growth, if *Q >* 0.

We have illustrated this numerically with model simulations. When comparing different multi-species systems in their resistance to invasion, for fixed parameters, we find that the higher the value of *Q*, the harder the invasion by outsider species (Fig. A-G). In contrast, we find no significant role of system diversity (Shannon, Gini-Simpson indices Fig.I-K), but a positive effect of the number of species for invader success (Fig. H). The final outcome of any given invasion however, as captured in Eq.4, besides system *Q*, depends also on other details of resident dynamics, and crucially, on the traits of the invader, relative to the resident community, at the invasion time. Notice that this does not necessarily imply restrictions on cooperation or competition as suggested previously (45), but rather a far more general condition about similarity or relatedness: the mean invasion fitness of the ‘invader’ must exceed the mean invasion fitness of the ‘resident system’.

### Invading the same system at different times: variability of outcome depends on mode of species coexistence

A corollary of this mathematical understanding of invasion is that following the dynamics of systemic mean invasion fitness 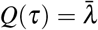, the timing of invasion (by the same invader) will likely have a different success and effect on a given system, depending on the actual magnitude of *Q*. We performed simulations of invasion in two qualitatively different systems to illustrate this phenomenon: i) in a system where multi-species coexistence tends to a stable fixed point (Fig.5A-B), and ii) in a system where underlying species coexistence is oscillatory (Fig.5G-H). In Fig. 5C-D, we show one illustration of dynamics of a particular successful invader in multispecies system 1 (invader colored in brown), and the corresponding dynamics of systemic colonization resistance *Q* after invasion. In Fig. 5I-J, we show one illustration of dynamics of another particular successful invader in multispecies system 2 (invader again colored in brown), and the corresponding dynamics of systemic colonization resistance *Q* after invasion.

**Figure 5.**
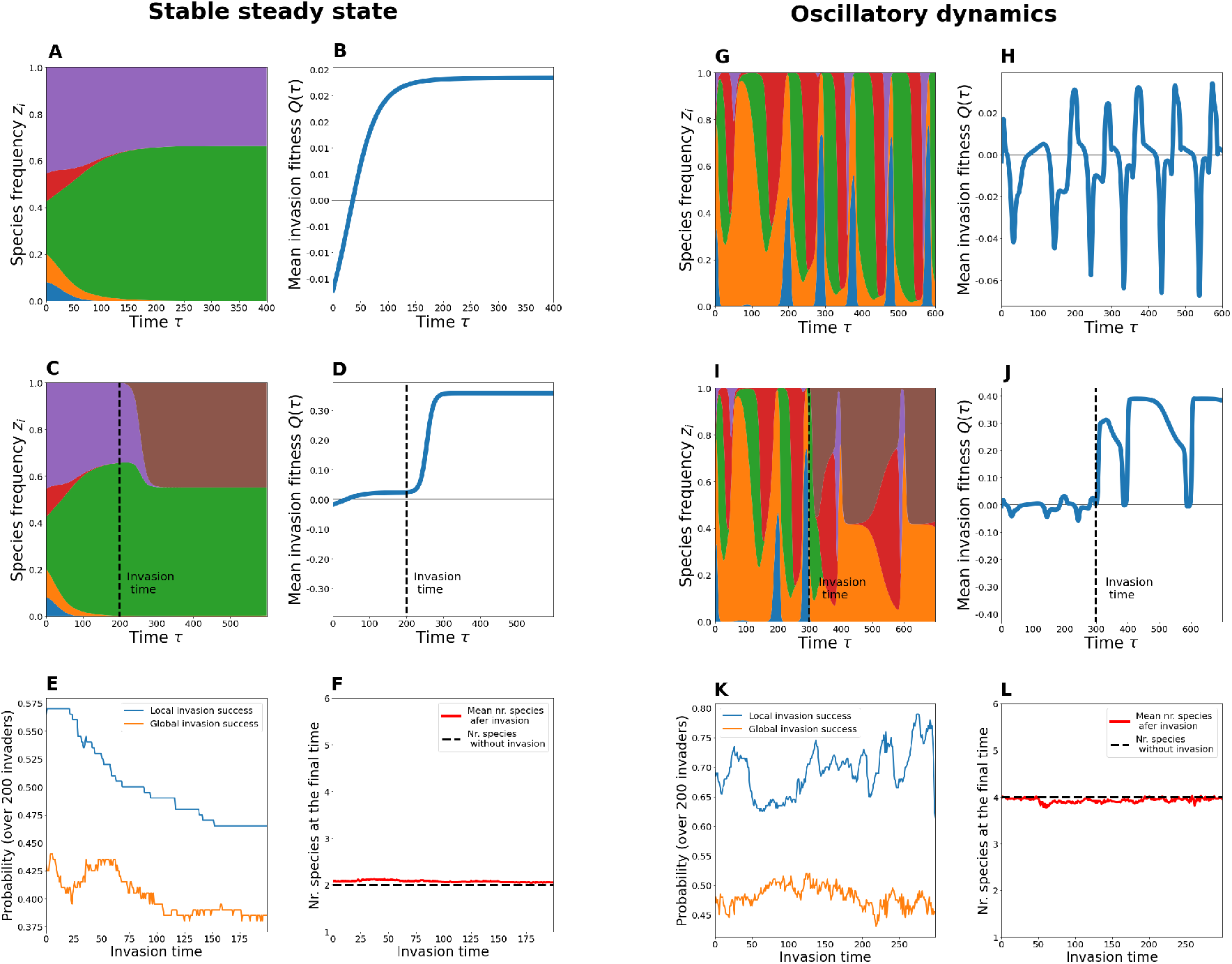
Two qualitatively different resident systems and sensitivity of invasion outcome to invasion timing. Here we fix two resident systems with *N* = 5 (A-F and G-L panels) and study the effect of invasion time over 200 random invaders. **A**., **G**. The baseline dynamics of species frequencies in the absence of invasion (system with stable steady state and oscillatory dynamics, respectively). **B**., **H**. The dynamics of mean invasion fitness of each system in the absence of invasion. **C**., **I**. Example of species frequencies for a particular invader (in brown) and a particular invasion time (dashed line). **D**., **J**. Corresponding mean invasion fitness dynamics for each system with invasion shown in (C) and (I). **E**., **K**. Summary of many invasion outcomes as a function of invasion timing in the same system. We sample 200 invaders for each time point, simulate invasion starting from low initial frequency of 1/%, and compute the proportion that results in initial growth (local invasion) and probability of ultimate persistence in the system (global success) using a dynamic horizon of *T* = 1000. In a system with monotonic 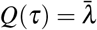, mean invader success tends to a constant, whereas in the system with oscillatory 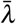 invader success tends to oscillate. **F**., **L**. One aspect of system outcome with and without invasion is the final number of coexisting species (here calculated as those above a given relative abundance threshold *z*_*i*_ *>* 1% during the last 10% of the simulation time interval, and averaged over many different simulations). On the left, we observe that invasion may increase slightly number of coexisting species if invasion happens early in the course of community dynamics, when the system is far from equilibrium. If invasion happens closer to equilibrium, the qualitative number of final coexisting species tends to be the same, but with one of the species effectively replaced by the invader. On the right, we observe that invasion on average reduces diversity slightly below its originally expected level.

Then we repeat such invasion experiments over different invasion time points, and multiple random invaders, and summarize the results. First, we confirm that temporal success of outsider invasion on average follows clearly the dynamics of *Q* of the underlying system: it tends to a constant when *Q* is monotonic (Fig.5E), and oscillates periodically when *Q* is periodic (Fig.5K). In particular, the less *Q* changes (i.e. when the underlying system tends to stable coexistence fixed points), the weaker the timing effect, except for a short transient whenever the resident community may be far from its equilibrium (Fig.5E), and it may seem that the probability of successful invasion decreases with invasion time, until it saturates. Conversely, the more oscillatory colonization resistance *Q* is (i.e. the system is characterized by limit cycles), the stronger the timing effect on average, and invasion success can vary more along time (Fig.5K), albeit being overall relatively higher.

In general, these patterns persist both when computed locally (invader growth after invasion) and when computed globally (invader persistence after invasion), with the relative difference between the two probabilities being about 20%. This suggests that on average the final fate of invaders may be quite predictable based on their initial invasion dynamics: if they invade, most likely they will persist (70-80% of invaders that do experience some growth in the system, will persist).

Our simulations also suggest that mean invasion fitness tends to increase after invasion (Fig.5D,J), a result that is plausible when considering that a pre-requisite for successful invasion is 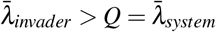, so if the invader eventually manages to stay in the system it does so via or with augmenting systemic *Q*. In the stable coexistence case, the successful invader tends to replace one species in a given niche - thereby not affecting final number of coexisting species *n* (Fig.5F), but increasing the stability of this coexistence. In the oscillating case, such increase in coexistence stability comes from the successful invader more often decreasing the number of coexisting species (Fig.5G).

In fact, the further from equilibrium the system is at the moment of invasion (the earlier the invasion relative to a system’s development), the more likely it is that the invader disrupts final outcome, and viceversa; the closer to equilibrium, the less the invader can perturb the system. In particular, in the stable coexistence case, when considering the number of coexisting species after invasion, we find it can reach as high as 5 species if the invasion happens early, in a system where it is expected to be 2 under its natural course, but on average, tends to equalize that of the original system without invasion for later timings (Fig.5F). Whereas, in the oscillatory case, successful invasion typically results in a slight average decrease of the number of species below the baseline expected with the original system (Fig.5L), having either way an overall stabilizing effect on coexistence.

### Invading different systems: success varies with invasion architectures of resident species

We finally studied random invader success in given community contexts, fixing invasion time, but now varying systematically the underlying mutual invasion matrix Λ of the resident species (Figure 6). We performed 50 simulations of random outsider invasion, for 200 realizations of each canonical Λ structure: (invader-driven, resident-driven, symmetric, anti-symmetric and random), and computed two indicators of successful invasion: i) the probability of initial invader growth (local invasion); and ii) the probability of final invader persistence in the system (global invasion).

**Figure 6.**
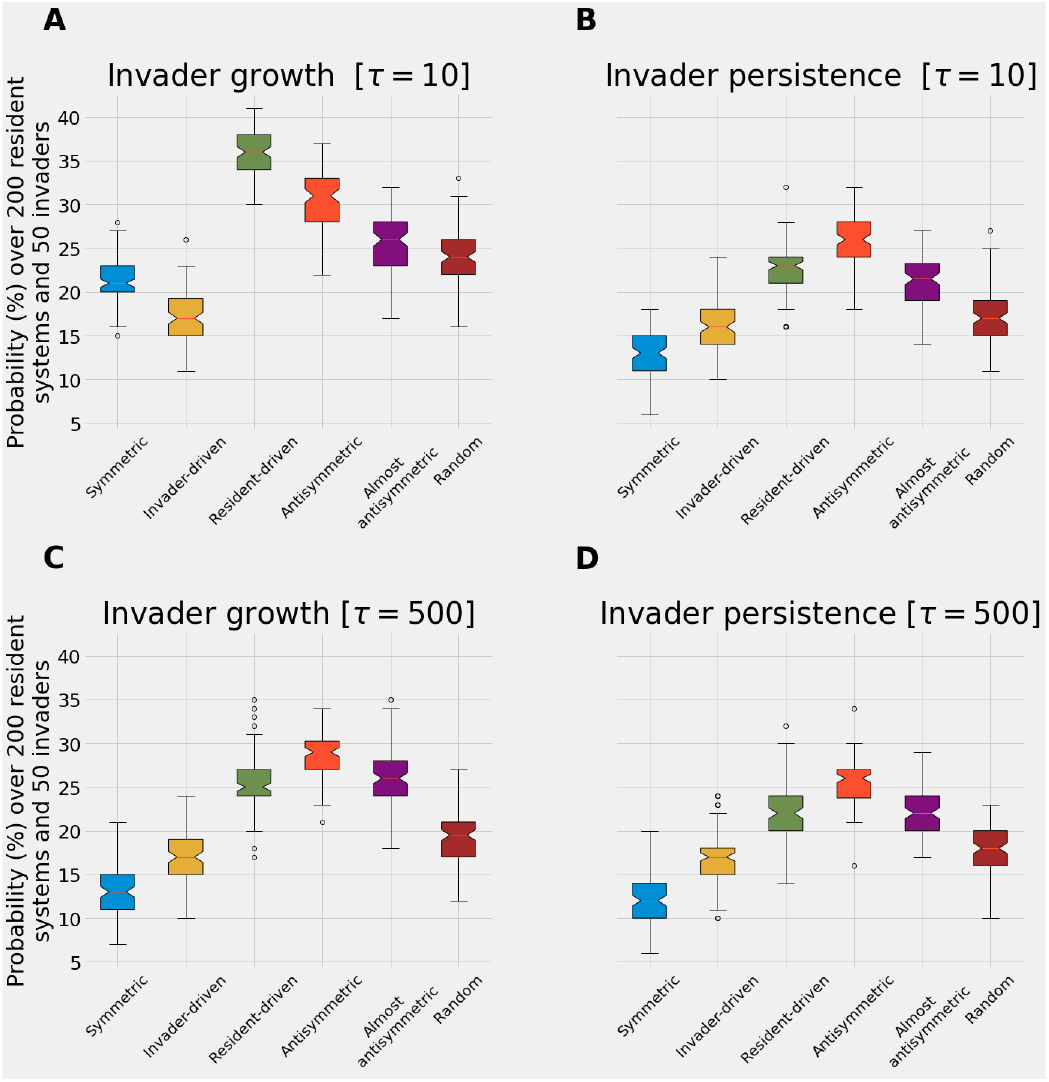
Mutual invasion structure of the resident microbial consortium and random invader success. We generated 200 random Λ matrix structures of size *N* = 5, with entry coefficients in the range [−1, 1], for each of 6 canonical architectures: symmetric, invader-driven, resident-driven, anti-symmetric, almost anti-symmetric, and random. For each random invader (random uniform 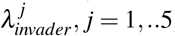), we simulated invasion in 200 systems of each type (Λ matrices), starting at the same initial frequency of the invader of 1% and equal relative frequencies of the resident 5 species totalling 99%. Over 200 systems, we computed the probability of outcome for each invader, summarized in the colored boxplots, which show entire variation over 50 invaders for each type of Λ matrix. Systems were run for a time horizon of *T* = 1000. The invader was introduced at two timepoints: (**A-B**) early in the development of the system close to initial equiabundance (*τ* = 10), and (**C-D**) later in the development of the system, closer to its intrinsic equilibrium (*τ* = 500), which contains more signature of Λ-driven selection. Local invasion counts all those instances where the invader grew from its initial frequency at some point from introduction up to *T* = 1000. Global invasion counts all those instances where the random invader is still present in the system at final time *T* = 1000. The ranking observed between Λ structures can be very useful when engineering multi-species communities for the purpose of maximizing colonization resistance.

As expected from our previous analysis (18) and the dynamics of *Q*(*τ*) in Figure 3, we find that the mutual invasion structure in the resident community strongly impacts invader success. Importantly, the highest probability of invasion, both local and global, is for the anti-symmetric invasion architecture of the resident species, yielding *Q* = 0 as the most favourable invasion context for outsiders. In contrast, the lowest probability of invader growth and invader persistence is found for symmetric, invader-driven and random mutual invasion of the resident species, indicating that these communities contain the least favourable (i.e. highest *Q*) colonization resistance for outsider invasion. This ranking is robust to random variation in the resident (200 different realizations of the same structure) and invader invasion coefficients (50 different invaders), pointing to a key signature of system stability and invasibility encoded precisely in the Λ matrix type. Although if introduced early (*τ* = 10), the probability of outsider species initial growth is lowest in a community characterized by invader-driven (row-wise variation) Λ architecture (Figure 6A), ultimate outsider persistence in the system (Figure 6B), and in general both outsider growth and persistence for later invasions are minimized if the underlying system displays symmetric mutual invasion structure (blue boxplot in Figure 6C-D).

Crucially, the ranking of these mutual invasion structures (Λ matrix types) suggests that the most resistant, externally stable communities are those whose constituent species ‘play’ symmetric or hierarchical invader-driven invasion ‘games’. Such Λ matrices, in our trait-based model, may be both obtained via special cases of symmetric co-colonization susceptibilities *K*_*i j*_ (see Table 2 in (18)). This suggests that stronger collective coexistence may indeed arise when species derive symmetric benefits from shared resources. Ultimately, the type of multispecies dynamics critically depends on the corresponding mutual invasion matrix. Hence, even though the 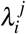 may be derived from particular biological traits and processes, which are of course model- and system-dependent (20), multispecies dynamics, evolution and composition over time, will eventually come down to the net Λ matrix and its effective adaptation under replicator dynamics.

While we could not explore all the complexities of the interplay between species traits and systemic colonization resistance, the results presented here should provide a solid ground for deeper analytical advances and mechanistic hypothesis-driven empirical investigation. We highlight that the key to study system invasibility is precisely the architecture and magnitude of mutual invasion coefficients among constituent species.

## Discussion

Colonization resistance in microbiology (2) is closely linked with the theory of invasibility in ecology (46; 47; 48), long studied but rarely integrated between the disciplines. In the late fifties, Elton argued that complex communities should be more resistant to invasion by new species (49). Later May (50; 51) showed that complex ecological communities tend to be less stable, i.e. in these systems it is harder to return to equilibria from small perturbations to existing species. Since then, the complexity-stability debate began, inspiring many investigations (46). Crucial to study invasibility and stability of ecosystems have been simulations, using random interaction networks or varying their properties, e.g. the percentage of cooperative vs. competitive links (52), or examining the relative interaction in invader vs. within-resident community (45). More recently, theoretical approaches to ecological invasion are also invoking trait-based invasion fitness concepts and characterizing the trait distance between resident and invader species for successful invasion (53).

However until now, to our knowledge, no unifying formal analytical quantity has emerged for colonization resistance per se, that integrates species interactions and traits and resource availability. Two notions of stability of a multispecies system, internal vs. external, have been studied, with the latter receiving much less attention. It is external stability that relates most with colonization resistance. While the stabilizing vs. destabilizing role of mutualism and cooperative interactions among species, is still theoretically debated (22; 52), as are diversity effects on external stability (54), recent work is attempting to reconcile Elton’s and May’s views (55) and seeking deeper and concise mechanistic insight.

In this report, we propose a fresh conceptual framework to bridge between these fields and contribute to the complexity-stability debate, by calling attention on the thus-far neglected measure of mutual invasion fitness between any two species, 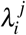, and the various topologies of the matrix it generates. This higher-level quantity embeds within it any lower-level traits and asymmetries, including cooperation/competition (in our case *K*_*ij*_), and mean-field resource availability indicators (in our case *R*_0_, *k*), and evolves in its own right following species frequency dynamics, as a collective feature of the system. With this framework, we propose unification between ecological, evolutionary dynamics and emergent system properties in multi-species communities, in the same spirit of reconciling simplicity with complexity in functional biology, advocated by others (56), and of improving predictability of microbial ecology models (57). Our derived replicator equation for multi-species dynamics, obtained from an analogy with an epidemiological model, simplifies complexity, while integrating cooperation and competition in the same continuum via the parameter *k*, naturally linking with Lotka-Volterra models (23), putting the concept of invasion fitness from adaptive dynamics (24) at its core, and relating with the broad ecological theory of invasibility (46; 47; 48; 58).

The whole-community behavior exhibited in our model is a generic consequence of dynamic resource availability, similar to (59), where, in our case, abstract resources are amplified or diminished in value, by the simple co-presence of other ecological partners. In our closed system, essentially, species are simultaneously the ‘active’ entities, striving to survive, and the very ‘substrates’ upon which their life depends. There is no requirement for cooperative interactions in order for *Q* to evolve as a community-level function; only the possibility of species micro-scale co-occurrence, *k >* 0, is needed, coupled with the assumption that organisms transform their environment in slightly different ways. Then it becomes another application of the basic notion that fitness is context-dependent (21; 60; 61; 62).

Many theoretical phenomena, like those explored above and beyond, directly accessible via this formalism, could be empirically matched when comparing gradients in microbiota dynamics and composition, caused by antibiotics, diet, or other stressors in multispecies communities, and relating such gradients to mathematical instances of classical replicator dynamics (19; 43). Empirically these gradients could include variation across human or mammalian hosts of different age or conditions, or environments under the effect of nutrients and abiotic factors such as moisture, light, temperature (e.g. microbial consortia in soil, cheese rinds, natural biofilms). With model extensions under way (20; 39), to increase scope and generality, for example more trait dimensions for species differences, including growth and clearance biases besides interaction coefficients, possibly relevant to more biological scenarios (63), we show the backbone of the replicator equation formalism, in terms of mutual invasion fitnesses, stays the same. This constitutes a big advantage as theoretical study of replicator dynamics has a long history and solid mathematical foundations (21). In our view, the precise notion of colonization resistance, derived from this model as mean invasion fitness of a community, can enable deeper study of the role of species interactions on the ecology of antibiotic resistance (64), of general community-level cohesion independently of cooperation or competition (59), and provide more precise quantifiable links with data, both from *in-vitro* and *in-vivo* microbial ecosystems under evolution and adaptation (9; 36; 65). Furthermore, beyond colonization resistance, due to the explicit nature of the dynamical equations for multi-species frequencies, other traits of interest in the system could potentially be tracked over time, as a direct signature of ongoing selection.

In light of increasing calls for models to better describe microbial consortia and predict system-level properties (56; 57; 66), here we outline a new path for explicit mathematical investigation of colonization resistance and mean fitness in a microbial ecosystem. This framework may help synthesize a new understanding of the ecology, development, and evolution of host health and resilience to disease, mediated by microbiota.

## Methods

### Model for *N*-species varying only in co-colonization interactions *K*_*i j*_

In the model for *N* species, described in detail in (18) we track the variables *S, I*_*i*_ and *I*_*i j*_ that refer to the proportion of uncolonized niches, those singly-colonized by species *i* and those co-colonized by species *i* and *j*. The dynamics of colonization and co-colonization are given by the following equations:

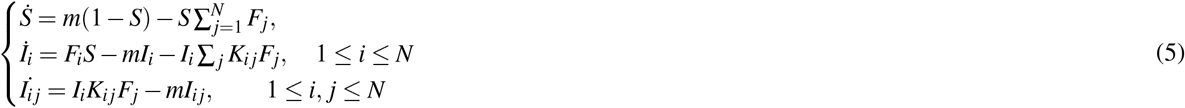

where 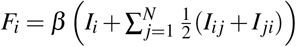 gives the force of propagation of species *i* in the system, summing the contribution of all micro-niches emitting propagules of that species. Because 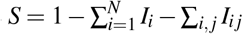, the dimension of the system is effectively *N* + *N*(*N* −1)*/*2. In this model in micro-niches co-colonized with different species *i* and *j*, the opportunity of each species to grow and be propagated is equal, thus represented with the 1/2 probability. In the system, the composite parameter *m* = *γ* + *r*, represents the microbial population turnover rate, encapsulating both the clearance rate *γ* of colonization episodes and the recruitment rate of free niches *r*. It is assumed that natural replenishment is balanced by natural mortality rate *r* = *d*.

The way in which each species interacts with itself and other species in the system upon co-colonization, is represented by the matrix *K*, where values *K*_*ij*_ above 1 indicate pairwise facilitation, while values of *K*_*i j*_ below 1 reflect inhibition between propagules of *i* and *j*, and where the order of colonizer and co-colonizer matters. In (18), we derive a slow-fast timescale decomposition of the dynamics, assuming similarity among species, in our case, in co-colonization interaction coefficients. Mathematically this is obtained by writing every *K*_*ij*_ as: *K*_*i j*_ = *k* + *εα*_*ij*_ under the assumption that *ε* is small, and *k* is a benchmark, e.g. the mean of 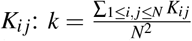. This leads to a normalized interaction matrix 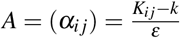 with the same distribution as *K*, but with mean 0 and variance 1 (if *ε* is taken to be the standard deviation of *K*), at the start of dynamics. With this formulation, the *N*-species frequency dynamics over the timescale *εt* are captured by the replicator equation 1. For more details, on the link with the mutual invasion fitness matrix Λ and global mean-field parameters, such as *R*_0_ and *k*, see Table 1 and (18; 19). For a more step-by-step intuition on the analogy we develop here to apply the SIS model framework to microbiota dynamics see Table S1.

### Generalized model for *N*-species varying in *K*_*ij*_ and 4 additional traits also satisfies replicator dynamics

The more general model for variation in other trait dimensions between similar species can be found in (20) where we formally derive that a similar replicator equation in terms of invasion fitnesses regulates the dynamics, but the mutual invasion fitness expression 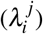 depends now on *all* traits. Besides the co-colonization interaction matrix *K* = (*K*_*ij*_), describing micro-scale susceptibilities to co-colonization (varied here) the 4 new trait axes we have included and studied in (20; 39) are: i) species-specific transmission/proliferation rates *β*_*i*_, ii) species-specific clearance rates in single colonization *γ*_*i*_, iii) pair-specific clearance rates in co-colonization (*γ*_*ij*_), and iv) species onward propagation biases from mixed co-colonization, depending on their order of arrival, 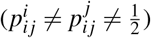. In this more complex scenario, the definition of ‘global context’ changes, as it may comprise also mean-field parameters in other traits, besides *R*_0_, *k* (relevant for the *K*_*ij*_-only model used here). The two-species system, namely the *N* = 2 dynamics in the case of 5-dimensional trait variation, is studied in (39), where the effect of the single to co-colonization ratio *µ* is shown to be very important, but this time influencing 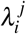 in a non-linear fashion. The dynamic colonization resistance of the *N*-species community emerges in this model again as a mean invasion fitness over all pairs in the system 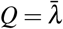, dynamically shaped by their changing frequencies. This model generalization shows that studying system dynamics and evolution in terms of the mutual invasion coefficients between constituent species is robust and widely-applicable.

## Author contributions statement

E.G. conceived the study, E.G. and S.M. conducted the investigation. E.G. wrote and revised the manuscript with critical feedback at all stages from S.M.

## Acknowledgements

The work benefitted from PESSOA grant 5666/44637YE awarded by Portuguese Foundation for Science and Technology (FCT) (to EG and SM), and cooperative agreement 2020-2001-230-Y20V7 by Le Studium, Loire Valley Institute of Advanced Studies (EG). Erida Gjini acknowledges support also by CEECIND/03051/2018. The format for this preprint is adapted from the Scientific Reports template available on Overleaf.com.

## Additional information

**Some codes for model simulation can be found on [GitHub](https://github.com/stenmadec/Replicator-equation-for-co-colonization/blob/main/Invasion_in_replicator.ipynb)**;

**The authors declare no competing interests**.

## Notes

### Competing Interest Statement

The authors have declared no competing interest.

