## Supplementary Movie Captions and Table S1 for "Towards a mathematical understanding of colonization resistance in multispecies microbial communities"

### (Movie file captions)

**Movie S1** **Illustration of model dynamics in co-colonization and species frequency space (example 1).** Here we visualize multi-species dynamics and evolution of system invasibility (Madec and Gjini, 2020 [1]), following the derived replicator equation 1. We used randomly generated $K_{ij}$, $\mu = 0.5$, $k = 1$, $\varepsilon = 0.1$, $\beta = 2$, and random initial conditions among $N = 10$ species. **A**. Pairwise invasion network structure. In **B**, we show the evolution of relative abundances over time. In **C**, the width of the nodes is scaled according to the frequency of each species ($z_i$), the width of the edges between any two species is scaled according to their product ($z_i z_j$). **D** Mean invasion fitness $Q$ displays mainly positive values over time, indicating high resistance to invasion of this system.

**Movie S2** **Illustration of model dynamics in co-colonization and species frequency space (example 2).** Here we visualize multi-species dynamics and evolution of system mean invasibility, following the replicator dynamics in Equation 1, with another set of randomly generated $K_{ij}$, $\mu = 0.5$, $k = 1$, $\varepsilon = 0.1$, $\beta = 2$ and random initial conditions among $N = 10$ species. **A**. Pairwise invasion network structure. In **B**, we show the evolution of relative abundances over time. In **C**, the width of the nodes is scaled according to the frequency of each species ($z_i$), the width of the edges between any two species is scaled according to their product ($z_i z_j$). **D** Colonization resistance $Q$ displays wild oscillations over time, spanning the positive and negative range, indicating transient windows of time where this system favours outsider species invasion.

**Movie S3** **Increasing basic reproduction number $R_0$ has system-specific effects on the outcome between two interacting species.** Here we visualize the context-dependence of species outcomes for $N = 2$, via $\lambda_i^j$ as a function of basic reproduction number $R_0$ (scaling the overall prevalence of colonization according to $1 - 1/R_0$ and the ratio of single to co-colonization $\mu = 1/(k(R_0 - 1))$). We compare two systems (1 and 2) which vary in primary $K_{ij}$ co-colonization interactions, but display same equilibrium frequencies $(z_1, z_2)$ between two species coexisting at $R_0 = 2$ (magenta line). Increasing $R_0$ has very different effects on the two systems. While system 1 (less competitive, higher $k$) preserves coexistence, system 2 (more competitive, lower $k$), for the same changes in $R_0$ shifts towards exclusion. This shows that perturbation effects depend explicitly on underlying species network. The rescaled interaction coefficients are as follows, for system 1: $\alpha_{11} = 0.2801, \alpha_{12} = 1.1202, \alpha_{21} = 1.4003, \alpha_{22} = 0.8402$ and for system 2: $\alpha_{11} = -0.9638, \alpha_{12} = 0.5507, \alpha_{21} = 0.6884, \alpha_{22} = 1.5145$.

**Movie S4** **Decreasing basic reproduction number $R_0$ has system-specific effects on the outcome between 2 interacting species.** Here we visualize the context-dependence of species mutual invasion fitnesses (ecological outcome) for $N = 2$, decreasing net microbial growth intensity $R_0$, hence reducing the overall prevalence of colonization according to $1 - 1/R_0$. We compare two systems (1 and 2) which vary in $K_{ij}$ co-colonization interactions (system 1 is less competitive than system 2), but display same coexistence between two species at $R_0 = 2$ (magenta line). Decreasing $R_0$ has very different effects on the two systems. While system 1 preserves coexistence for longer, system 2, for the same changes in $R_0$ shifts faster towards exclusion. This highlights that perturbation effects depend explicitly underlying species network. The rescaled interaction coefficients are as in Movie S3.

[1] Madec, Sten, and Erida Gjini. "Predicting n-strain coexistence from co-colonization interactions: epidemiology meets ecology and the replicator equation." Bulletin of mathematical biology 82.11 (2020): 1-26.

**Table S1. A step-by-step model derivation via Resource-Volume-Microorganism dynamics.** To relate the model to more classical and well-known chemostat dynamics models, we provide this possible illustration for how the dynamics can be 'seen' in terms of explicit fluxes between abstract 'volume' compartments. In the system without microorganisms, there is just resource ($R$) going into the system with rate $d$, equal to washout rate $d$, hence resource distribution and mixing in the 'volume' ($V$) of interest. The function $g(R)$ is a generic function describing how the resource spreads and binds to each unit of volume (micro-scale). The proportion of the volume that is 'free' of resource is denoted by $V_{free}$ and the proportion of the volume that contains resource is denoted by $V_R$. In the system with microorganisms, the relevant volume units for micro-organism growth become only the ones with resource, hence we have an expansion of $V_R$ into three sub-compartments: those without micro-organisms, those singly-colonized, and those co-colonized by microbial propagules. Assuming growth and release of propagules is instantaneous, we have that total number of microbial propagules in the system is a direct linear function of $V_{RI}$ and $V_{RD}$, with 'conversion rate' $\beta$. Mass-action kinetics and homogeneous mixing determine then transmission dynamics. The clearance rate $\gamma$ can be seen to encapsulate those processes by which the 'substrate-microbe' units can lose the proliferating microbes without losing the resource (death of bacteria, nutrient recycling..). Focusing on the sub-system with resource only, and rescaling all variables by their total sum, we obtain the variables $S, I, D$ in the core model structure proposed in the paper. Subsequently one can expand such structure to accommodate higher resolution in the parameter $k$, or other traits, to enable species-specific values and hence model compartments.

**System without microorganisms:** resource distribution and mixing

$$\frac{dR}{dt} = dR_{in} - dR \quad \text{(resource dynamics flux)}$$

$$\frac{dV_{free}}{dt} = -g(R)V_{free} + dV_R \quad \text{(volume units without resource)}$$

$$\frac{dV_R}{dt} = g(R)V_{free} - dV_R \quad \text{(volume units with resource)}$$

Equilibrium: $R \rightarrow R_{in}$ and $V_R \rightarrow \frac{g(R_{in})}{d+g(R_{in})}$ and $V_{free} = 1 - V_R$.

**System with microorganisms:** transmission among *"substrate-volume"* micro-units

$$\frac{dR}{dt} = dR_{in} - dR$$

$$\frac{dV_{free}}{dt} = -g(R)V_{free} + d(V_{RS} + V_{RI} + V_{RD})$$

*Relevant units supporting growth of the micro-organisms:*

$$\frac{dV_{RS}}{dt} = g(R)V_{free} - dV_{RS} - V_{RS}X + \gamma(V_{RI} + V_{RD}) \quad \text{(susceptible)}$$

$$\frac{dV_{RI}}{dt} = V_{RS}X - kV_{RI}X - \gamma V_{RI} - dV_{RI} \quad \text{(singly-colonized)}$$

$$\frac{dV_{RD}}{dt} = kV_{RI}X - \gamma V_{RD} - dV_{RD} \quad \text{(co-colonized)}$$

$$X = \beta(V_{RI} + V_{RD}) \quad \text{(micro-organism propagules)}$$

Equilibrium: $R \rightarrow R_{in}$ and $V_R = V_{RS} + V_{RI} + V_{RD} \rightarrow \frac{g(R_{in})}{d+g(R_{in})}$ and $V_{free} = 1 - V_R$.
Defining

$$S = \frac{V_{RS}}{V_R}, \quad I = \frac{V_{RI}}{V_R} \text{ and } \quad D = \frac{V_{RD}}{V_R},$$

we obtain the core SIS transmission model in the paper, with co-colonization for $S(t), I(t), D(t)$, with $S + I + D = 1$. Depending on the type of 'resource' ($R$) and microbes involved, the parameters $\beta, \gamma, k$ can vary in a system-specific manner.
